# Contribution of image statistics and semantics in local vs. distributed EEG decoding of rapid serial visual presentation

**DOI:** 10.1101/2023.09.26.559617

**Authors:** Eric Lützow Holm, Diego Fernández Slezak, Enzo Tagliazucchi

## Abstract

Spatio-temporal patterns of evoked brain activity contain information that can be used to decode and categorize the semantic content of visual stimuli. This procedure can be biased by statistical regularities which can be independent from the concepts that are represented in the stimuli, prompting the need to dissociate between the contributions of image statistics and semantics to decoding accuracy. We trained machine learning models to distinguish between concepts included in the THINGS-EEG dataset using electroencephalography (EEG) data acquired during a rapid serial visual presentation protocol. After systematic univariate feature selection in the temporal and spatial domains, we constructed simple models based on local signals which superseded the accuracy of more complex classifiers based on distributed patterns of information. Simpler models were characterized by their sensitivity to biases in the statistics of visual stimuli, with some of them preserving their accuracy after random replacement of the training dataset while maintaining the overall statistics of the images. We conclude that model complexity impacts on the sensitivity to confounding factors regardless of performance; therefore, the choice of EEG features for semantic decoding should ideally be informed by the underlying neurobiological mechanisms.

## Introduction

Humans interact with a multitude of different objects each day, recognizing and generalizing complex patterns of sensory information to identify and categorize them in concepts based on their perceived meaning (Logothetis and Sheinberg 1996). Vision and language are key to these tasks, since they allow to identify objects from concrete or abstract representations, as well as to rely information about them to other human beings (Bonner and Epstein 2021). Both processes are instantiated in neural networks spanning specific cortical regions, yet they also vary across individuals, depending on their personal experience, cultural background, and several other variables, highlighting the need to develop robust experimental procedures that are sensitive to these variations (Kuwabara and Smith 2016, de Bruine, Vredeveldt et al. 2018). To address these issues, human studies of visual object recognition generally combine neuroimaging techniques such as magneto- or electroencephalography (MEG, EEG) (Hebart, Contier et al. 2023) or functional magnetic resonance imaging (fMRI) (O’Toole, Jiang et al. 2005, Huth, Nishimoto et al. 2012) with sensory stimulation paradigms designed to present images that are categorized according to their semantic content (Robinson, Quek et al. 2023).

Generally, experimental paradigms to study visual categorization present subjects with approximately one image per second (Carlson, Tovar et al. 2013, Cichy, Pantazis et al. 2014, Kaneshiro, Perreau Guimaraes et al. 2015). However, recent EEG studies have shown that it is possible to extract information from brain activity elicited by subliminal stimuli, i.e. stimuli presented below the threshold of conscious awareness. This paradigm, known as rapid serial visual presentation (RSVP) (Grootswagers, Robinson et al. 2019, Gifford, Dwivedi et al. 2022), allows to obtain responses to large collections of images in comparatively short times, as highlighted in a recent publication using the THINGS dataset (Grootswagers, Zhou et al. 2022). This dataset comprises over 25000 manually curated natural images, each presented during 50 milliseconds. In particular, the data published by Grootswagers and colleagues contains EEG responses to 22248 natural images spanning 1854 concepts, acquired from 50 human participants. Given the high dimensionality of the data obtained in a RSVP paradigm combined with EEG recordings, a major challenge consists of separating informative neural activity patterns from signals that are not directly related to the meaning of the stimuli, such as the statistical regularities present in natural images, among other potential sources of bias (Robinson, Quek et al. 2023).

While previous studies addressed how variations in image statistics (such as luminance, contrast, etc.) influence the decoding of stimuli based on neuroimaging data (Campbell and Robson 1968, Geisler 2008, Masarwa, Kreichman et al. 2022), the contribution of these confounding factors to image classification accuracy has been scarcely explored for the THINGS dataset (Harrison 2022). Moreover, even though different feature selection methods have been applied to classifiers trained to decode semantic categories from brain imaging data, the interaction between model optimization and confounding sources of information remains to be systematically addressed. In particular, improving model classification accuracy could result in the selection of features that are not directly sensitive to the semantic information present in the images but instead represent low-level image statistics, which might elicit stronger and more systematic responses in the EEG signals, thus leading to low classification errors. Moreover, another method of representation similarity analysis (RSA) has been applied to determine whether the distance between patterns of elicited brain activity covariates with the degree of semantic similarity between the stimuli (Kriegeskorte, Mur et al. 2008). However, in this case the results depend on the features selected for the pairwise comparison of EEG responses, which may be more or less sensitive to low-level image information that can present significant correlations with the semantic distance. Overall, it is difficult to determine whether the EEG responses obtained from a RSVP paradigm reflect the outcome of object detection and categorization, or are driven by low-level information which cannot be fully disentangled from the semantic information present in the images.

The first main objective of the present work is to investigate whether dimensionality reduction applied to EEG recordings can simplify and improve the performance of models trained to categorize images. Second, we aim to determine the robustness of models of different complexity against confounding factors present in the data. We hypothesized that simpler models based on local information (e.g. single channel vs. group of neighboring channels) would be more sensitive to low-level information associated with the statistics of the natural stimuli, while the opposite would occur for more complex models trained with distributed signals. We performed our analysis on the EEG responses to a RSVP paradigm with images included in the THINGS datasets (Grootswagers, Zhou et al. 2022). We first optimized models by selecting an optimal combination of individual EEG channels and frequency bands, resulting in univariate models optimized to maximize the classification performance. Next, we compared their performance to that of models without feature selection, comprising all 64 EEG electrodes across a broad range of frequencies. Finally, we implemented different approaches to estimate the contribution of the low-level statistics of natural images to the performance of the different models, under the hypothesis of a different contribution to local and simple univariate models compared to distributed and more complex models.

## Materials and methods

### Paradigm overview and THINGS dataset

The EEG data was obtained from a previously published article and database (Grootswagers, Zhou et al. 2022). The original code published with the data is available from the Open Science Framework (https://doi.org/10.17605/OSF.IO/HD6ZK). Stimuli were presented in two phases as described in the original publication, both following a RSVP paradigm (Grootswagers, Robinson et al. 2019), where each image was shown for 50 ms before onset of the next visual stimulus (see **Figure 1A** for an illustration of the RSVP paradigm). EEG signals were filtered using a Hamming windowed finite impulse response filter with 0.1 Hz highpass and 100 Hz lowpass filters, re-referenced to the average reference, and downsampled to 250 Hz. Epochs were created for each individual stimulus presentation ranging from −100 to 1000 ms relative to stimulus onset. Two participants were discarded from the validation phase and one from the main phase, due to the presence of artifacts, resulting in severely distorted signals as well as evoked response potentials.

**Figure 1.**
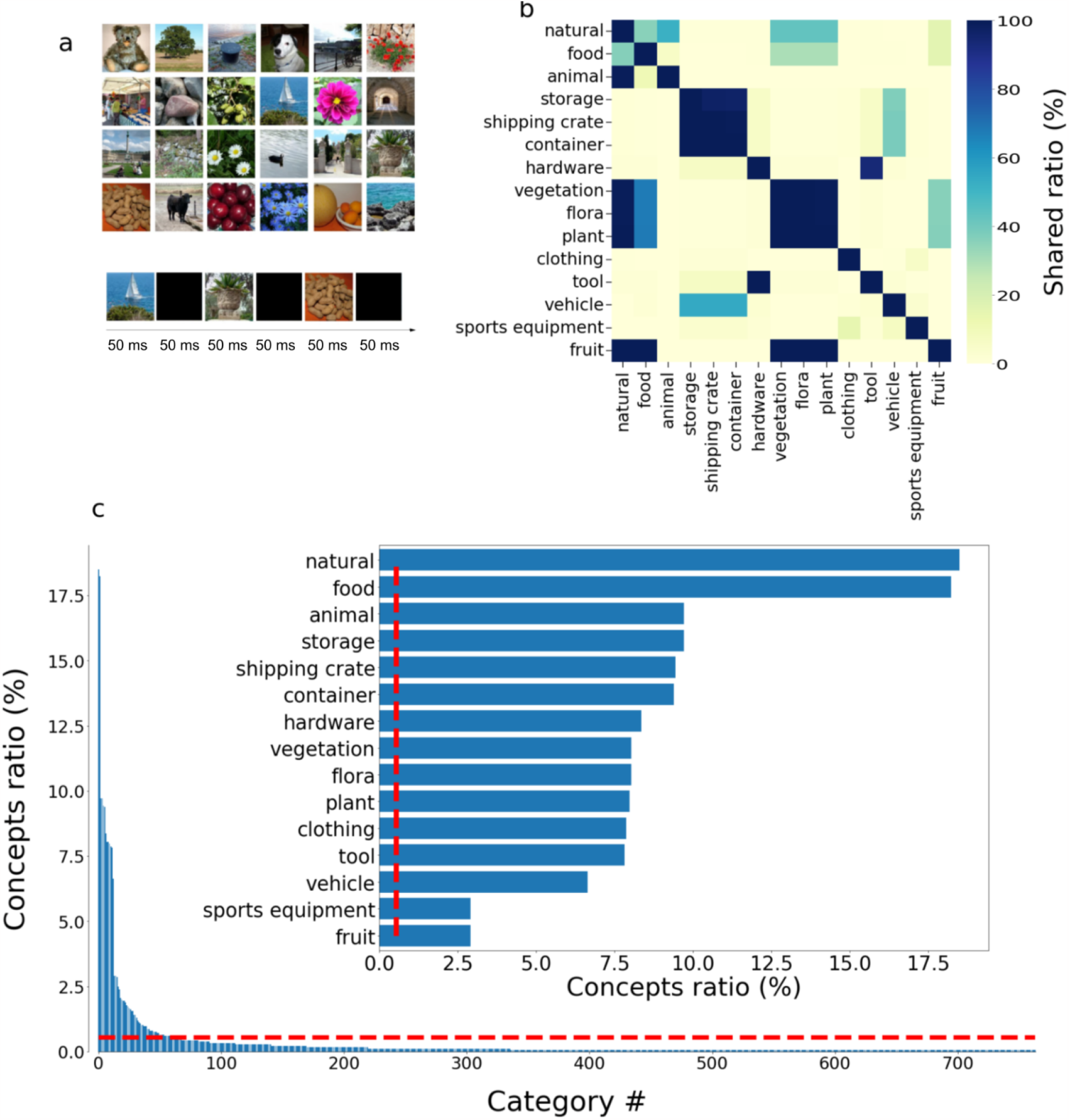
Outline of the THINGS dataset. A) Sample of randomly selected images in the dataset and outline of the RSVP paradigm, consisting of the presentation of an image for 50 milliseconds, with an empty screen lasting another 50 milliseconds before presentation of the next stimuli. B) Matrix representation of the number of shared images (as percentage of total) between the 15 largest categories in the dataset. While some categories are disjoint (e.g. “storage” vs. “animal”) others present a significant degree of overlap (e.g. “fruit “and “vegetation”). C) Ranking of categories based on the number of concepts they contain (as percentage of the total). The inset presents a detailed overview of the 15 largest categories. Red lines indicate the cutoff of 10 concepts per category.

The validation phase consisted of 200 images presented at randomized order, while the main phase consisted of 22248 natural images spanning 1854 concepts (12 images per concept), also presented in random order while avoiding the sequential presentation of two or more images belonging to the same concept. In turn, concepts could be grouped into categories according to their meaning. **Figure 1B** shows the 15 concepts with the largest number of concepts, as well as the overlap matrix between them. **Figure 1C** presents the ranking of all categories in the THINGS-EEG dataset based on the number of concepts they contain, as well as a detail of the top 15 categories in the inset.

### Linear discriminant analysis (LDA) model

The LDA model implemented a leave-one-out cross validation procedure combined with a regularized (λ = 0.01) linear discriminant classifier to distinguish between the images (validation phase) and concepts (main phase) (code implemented in scikit-learn, https://scikit-learn.org/). The cross-validation procedure yielded a mean accuracy value for each image or concept pair, participant and time sample. These accuracies can be understood as a dissimilarity metric, where a high value indicates that the images/concepts are dissimilar, thus facilitating their discrimination.

### Univariate feature selection

The feature selection procedure was based on computing the effect size (Cohen’s D) of the difference between the EEG signal corresponding to different image/concept pairs, and then obtaining the average across all pairs. Thus, high values of this metric indicate that, on average, the selected channel and frequency band present significant differences between concept pairs. This was repeated for each subject, EEG channel and time sample. After selecting the 10 highest ranking EEG channels according to this metric, we repeated the process across multiple band-pass filters within the range 0.5 to 100 Hz at 0.5 Hz intervals. Finally, the metric was averaged across participants.

### Model comparison

Different models were trained using the EEG data from either the validation or main phase of the experiment: 1) distributed (all EEG channels and broadband) model trained using regularized LDA; 2) local (single EEG channel and narrow band) model using the median of the EEG values obtained as response to presentations of two different images/categories, thus forcing a separation into two groups with the same number of elements; and 3) local (single EEG channel and narrow band) model trained using regularized LDA (same hyperparameters as in model 1). In particular, the computations underlying model 2 are very straightforward and more efficient in comparison to the other two models. To facilitate comparison between models, the corresponding performance metrics were adjusted based on their corresponding chance level, resulting in values ranging from 0 (random classification) to 1 (optimal classification).

### Natural image descriptors

The choice of image descriptors was guided by the previous report of Harrison (Harrison 2022) showing that better-than-chance classification can be obtained using only two metrics (luminance and luminance contrast). The list of 27 natural image descriptors reported by Harrison was enlarged to include the average value of the red, green and blue channels, resulting in 30 metrics. An iterative procedure was implemented to reduce the number of metrics relevant for image/concept classification based on the recorded EEG response. First, PCA was applied to the descriptors and the informativeness of the first principal component was assessed by computing its Pearson correlation vs. the voltage at the peak EEG response. Next, this procedure was repeated after subtracting one of the 30 metrics and determining which one led to the largest decrease in the Pearson correlation coefficient between the first principal component and the EEG response. Accordingly, one metric was eliminated after each step of the process, leading to a final subset of 8 image descriptors. To avoid over-fitting, the complete process was repeated 20 times using 33 random subjects for fitting and the remaining 16 for validation.

The final subset of descriptors comprised those which were not selected for elimination in any of the 20 repetitions, which were the following: luminance, average of the green and blue channels, two metrics related to orientation contrast energy and three related to the spatial frequency energy of the images. Luminance indicates the intensity of light from a given area of space after the light has passed through a model of the sensitivity of a standard eye (Harrison 2022). The orientation contrast energy is computed as the Fourier amplitude within each of 16 oriented bands spanning a full circle, obtained using oriented filters (Harrison 2022). This metric indicates whether the energy in the spectrum appears concentrated in specific directions. Finally, the spatial frequency energy was computed as the Fourier-transformed amplitude values, grouped according to their distance from the origin. This metric indicates whether the energy is concentrated at specific spatial frequencies.

### Clustering of images based on image descriptors

The 8 image descriptors previously found (see “*Natural image descriptors*”) were used as input to agglomerative hierarchical clustering (Ward’s method), yielding clusters (575, 762, 287 and 230 images each).

### Iterative replacement of concepts within categories based on image descriptors

A subset of the largest 37 concept categories was selected, spanning at least 1% of all concepts each. For each category, subject and time sample we computed the average Cohen’s D of the classifier performance scores (LDA and median) in response to concept pairs belonging to the same category, as well as for concept pairs with one member of the given category and the other belonging to a different one. Thus, a positive value of Cohen’s D implies that concepts from different categories are better distinguished compared to concepts within the same category, and vice-versa for negative values. Of all categories showing positive values, a subset of 4 categories was selected for further analysis (“human”, “body”, “animal” and “hardware”). Next, we selected a random concept in one of these categories and selected the nearest one that 1) belongs to a different category and 2) is the closest in terms of the 8 image descriptors described in the subsection “*Natural image descriptors*”. Finally, the concept was replaced and the effect size was re-computed. This process was repeated until all concepts were replaced, leading to semantically heterogeneous categories with concepts associated with homogeneous images in terms of their low-level descriptors. If the classifiers are sensitive to semantic information, it is expected that the effect size will gradually decrease, while the converse is expected if the classifiers are based on image statistics regardless of their perceived meaning. To estimate the effect of this random replacement, we used least-square linear fits to obtain the slope in the plot of change in effect size vs. % replacement, as well as the associated p-values.

## Results

We first performed a feature selection procedure to optimize model performance by exploring different choices of spectral content and EEG channels. For this purpose, we first analyzed the validation data published by Grootswagers et al., consisting of EEG responses to 200 images, each presented 12 times following a RSVP paradigm. In the original analysis by Grootswagers et al., linear discriminant analysis (LDA) was used to construct pairwise classifiers based on data from the 64 EEG channels at a wide range of frequencies (0.1 to 125 Hz); in the following we refer to these as models using distributed information (or distributed models). We attempted to reproduce the results in Grootswagers et al., but using simpler models comprising individual EEG channels at narrower frequency bands, which we refer to as local models. We note that this labeling is approximate, since individual electrodes receive information from multiple sources due to volume conduction effects; however, disentangling these sources from a single time series is not possible, whereas the use of information from all electrodes could in principle provide complementary information capable of capturing multiple sources of neuroelectric activity.

To investigate how EEG can distinguish between images with data gathered from a single channel, we grouped the 12 presentations of each image and analyzed the time lapse between -100 ms and +1000 ms around the presentation of each stimulus, which lasted for 50 ms. For each time sample within this range, we computed the average effect size for the difference between all pairs of images as a proxy of the average pairwise discriminative power of the local EEG signal across time. This procedure was repeated for each EEG sensor and each of the following frequency bands: 0.5-4 Hz, 4-8 Hz, 8-12.5 Hz, 12.5-30 Hz, 30-50 Hz, 50-70 Hz and 70-100 Hz (approximately corresponding to delta, theta, alpha, beta and gamma bands). The results of this analysis are shown in **Figure 2A**, where occipital and parietal channels (P3, P7, O1, Oz, O2, P4, POz, PO4, PO8, P2) appear highlighted the most informative to distinguish between image pairs. **Figure 2B** presents the average effect size across all channels as a function of time, with two clear peaks around 200 ms after presentation of the stimulus. **Figure 2C** presents the same information but in terms of EEG topographic maps, where the most informative channels appear marked using black dots, highlighting that most of them are located in the occipital region.

**Figure 2.**
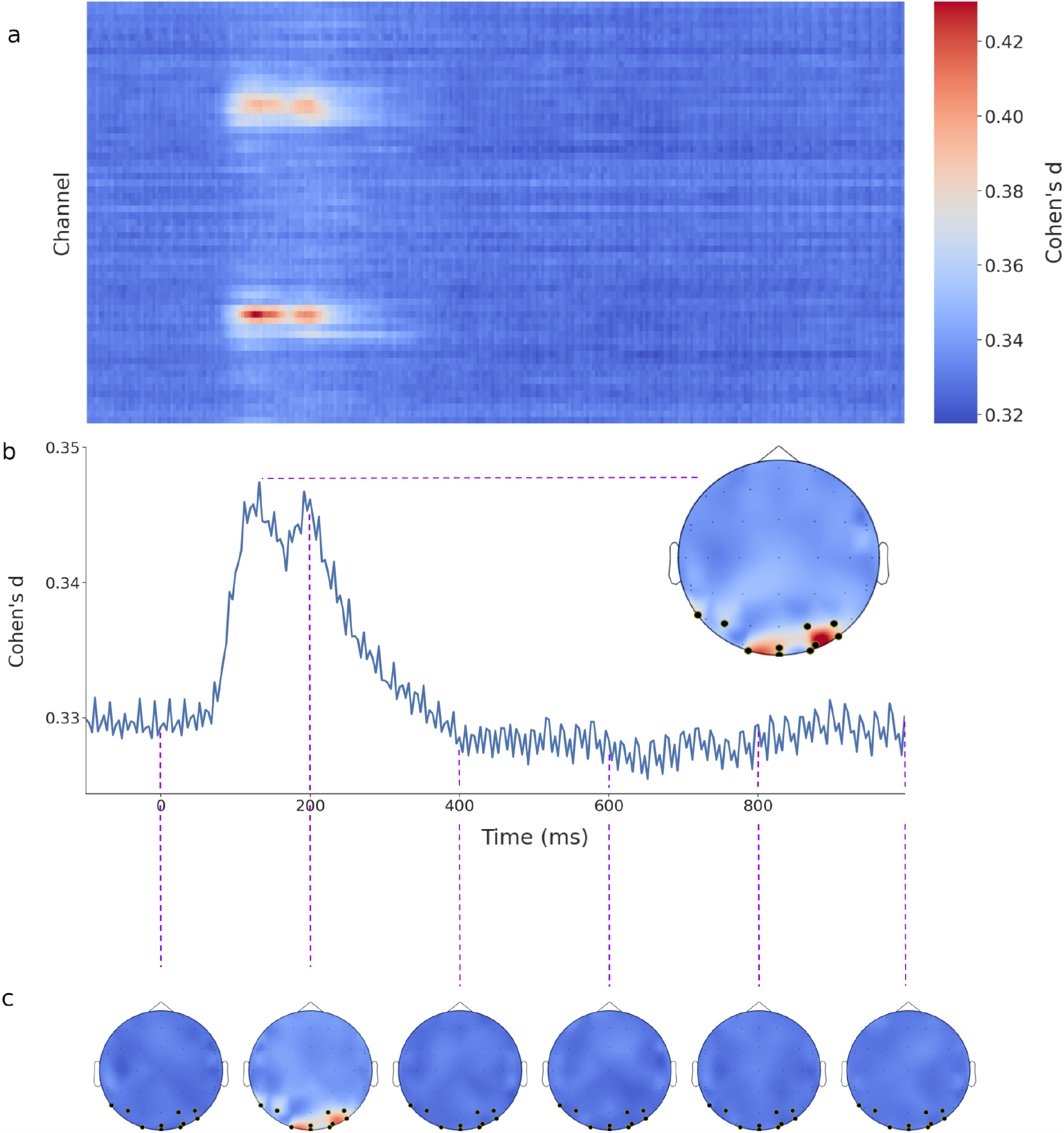
Informativeness of individual EEG channels to distinguish between pairs of images. A) Matrix representation of the average effect size (Cohen’s D) between all pairs of images per channel and time sample. Channels with high average effect size are located in occipital and parietal regions (P3, P7, O1, Oz, O2, P4, POz, PO8, P2). B) Average effect size across all electrode pairs, showing distinct peaks around 150 ms and 200 ms. Topographic maps present the scalp distribution of effect sizes at 100 ms intervals, while the inset highlights the 10 most informative channels at around 200 ms after stimulus onset. Overall, we observe that for this task the optimal channel and frequency combination is given by PO4 at the 4-12.5 Hz range (see Figure 3B for a comparison of results obtained using PO4 at different frequency ranges).

**Figure 3A** displays the effect size obtained from all the selected channels as a function of time in a single plot to facilitate their comparison, while **Figure 3B** shows this information for the optimal channel PO4 at different frequency bands, where it is clear that the broadband signal results in worse accuracy relative to narrower bands starting at 4 Hz. The simplest model that can be constructed based on this information uses the median of the EEG values as the threshold for classification, thus forcefully dividing the samples into two categories. **Figure 4C** presents a comparison of three different models: LDA trained using broadband data from 64 channels (Grootswagers et al., 2021), LDA trained using data from PO4 at 4-12.5 Hz, and the model which uses the median as threshold for classification. The last two models present similar accuracy values, and both are based on data from a single channel at a narrow frequency band.

**Figure 3.**
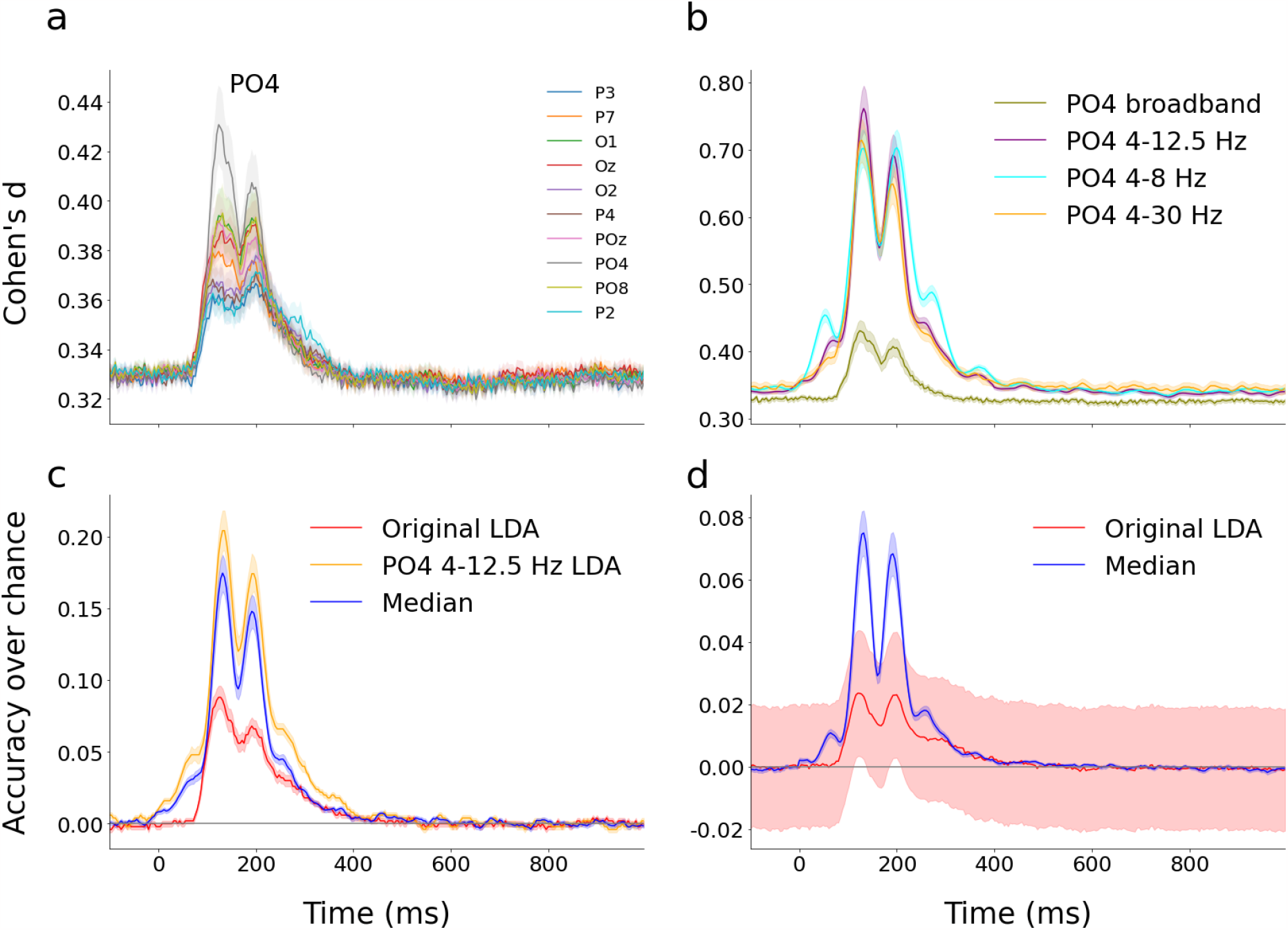
Comparison of models used to classify images in the THINGS-EEG validation data (panels A to C) and in the main phase of the experiment (panel D). A) Average effect size across image pairs for the 10 best channels. B) Same as in panel A but for channel PO4 at different frequency ranges. C) Comparison of the LDA distributed model based on 64 broadband EEG signals (Grootswagers & Robinson, 2021), the local LDA model based on PO4 at 4-12.5 Hz, and the local model obtained by separating using the median as classification threshold. D) Comparison of the distributed LDA model based on 64 broadband EEG signals (Grootswagers & Robinson, 2021) and the local model obtained using the median as classification threshold, for the main part of the THINGS-EEG dataset where models distinguish between pairs of 1854 concepts with 12 images each.

**Figure 4.**
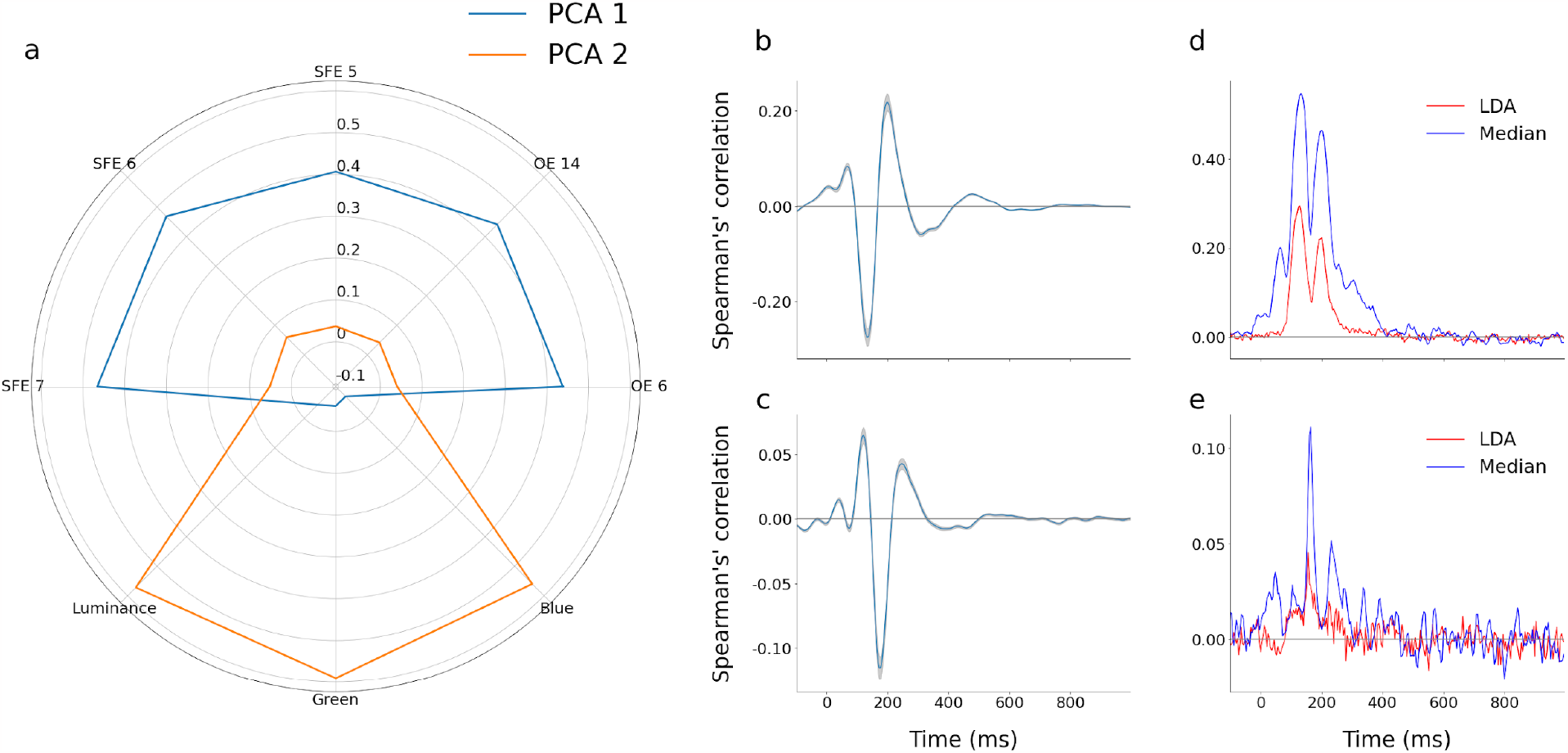
Explaining the EEG response and the classification accuracy in terms of systematic differences in the image statistics. A) Radar plot showing the composition of the two principal components (PCA 1 and 2) of a reduced set of 8 descriptors that quantify the statistics of the natural images presented to the participants. B) Spearman correlation coefficient between the first principal component value and the EEG signal at PO4. C) Same as in panel B, but for the second principal component. D) Correlation coefficient between the pairwise difference of the principal component averaged across all images in each concept, the accuracy of the distributed model using all EEG channels (LDA), and the local model based on splitting PO4 values by the median (median). E) Same as in panel D, but for the second principal component.

Next, we extrapolated this procedure to the main phase of the experiment reported in the THINGS-EEG dataset, comprising 1854 concepts with 12 images each (22248 images in total). Moreover, the concepts can be grouped into larger categories based on their meaning (e.g. “eye”, “arm”, “nose” could belong to a larger category termed “body”). Figure 3D presents a comparison of the multivariate LDA distributed model vs. the classification by the median using local information (PO4 in the 4-12.5 Hz range). Again, we can see that the latter model shows higher accuracy values at around 100 ms and 200 ms after stimulus onset.

After assessing the performance of these two models to distinguish between concepts in the THINGS-EEG dataset, we assessed their robustness against potential confounds stemming from regularities in the statistics of the natural images. For this purpose, we characterized each image with a series of quantitative descriptors (see Methods section for a detailed description). We first applied a cross validation procedure to select the top 8 image descriptors presenting the largest correlation with the PO4 EEG signals, resulting in the following set of descriptors: luminance, green and blue intensity, two descriptors related to orientation contrast energy (OE) and three related to spatial frequency energy (SPE). We then applied principal component analysis (PCA) to these 8 descriptors to further restrict the analysis to the two components explaining the largest variance of the images in the dataset (see **Figure 4A** for a radar plot representation of the principal components). **Figures 4B** and **4C** present the Spearman correlation coefficient of each of these components and the value of the EEG response at PO4, where two peaks (negative and positive, respectively) can be seen after 100 and 200 ms of the stimulus presentation. Finally, we tested whether these descriptors contribute to the classification of the image concepts. For this, we computed the average value of the first principal component for all images associated with each concept, and then for each pair of concepts we computed the difference between these averaged values. **Figure 4D** shows the Spearman correlation between these differences and the accuracy obtained using the distributed LDA model, as well as the local model using the median as threshold. In this figure, the size of the correlation coefficient indicates how much of the variance in the classification accuracy can be explained by differences in the image statistics. The model using local information (narrow band PO4) presented correlation coefficients considerably larger than those obtained using the distributed model (LDA using all EEG channels). **Figure 4E** presents the results of the same analysis, but using the second principal component instead; in this case, the correlation coefficients are overall smaller but still the local model still presents the largest values.

For further validation, we determined the concepts showing the largest average first/second principal component values as the first quartile of the distribution. We computed the effect size of the accuracy scores (obtained using the LDA and median models) between the concepts in the first quartile and the rest, with results shown in **Figure 5A** and **5B** for the first and second principal components, respectively. An effect of the image statistics is suggested by the comparatively large effect size, indicating that the first principal component of the image statistics descriptors systematically influenced the classification scores. Moreover, we observe that the effect size is higher for the local model (median) vs. distributed model (LDA), suggesting that the former is less robust against regularities in the natural images. **Figures 5C** and **5D** show the Spearman correlation between the dissimilarity vector (for each pair of concepts, the difference between the first and second principal component values) and the LDA and median scores, which contain similar information as panels A and B.

**Figure 5.**
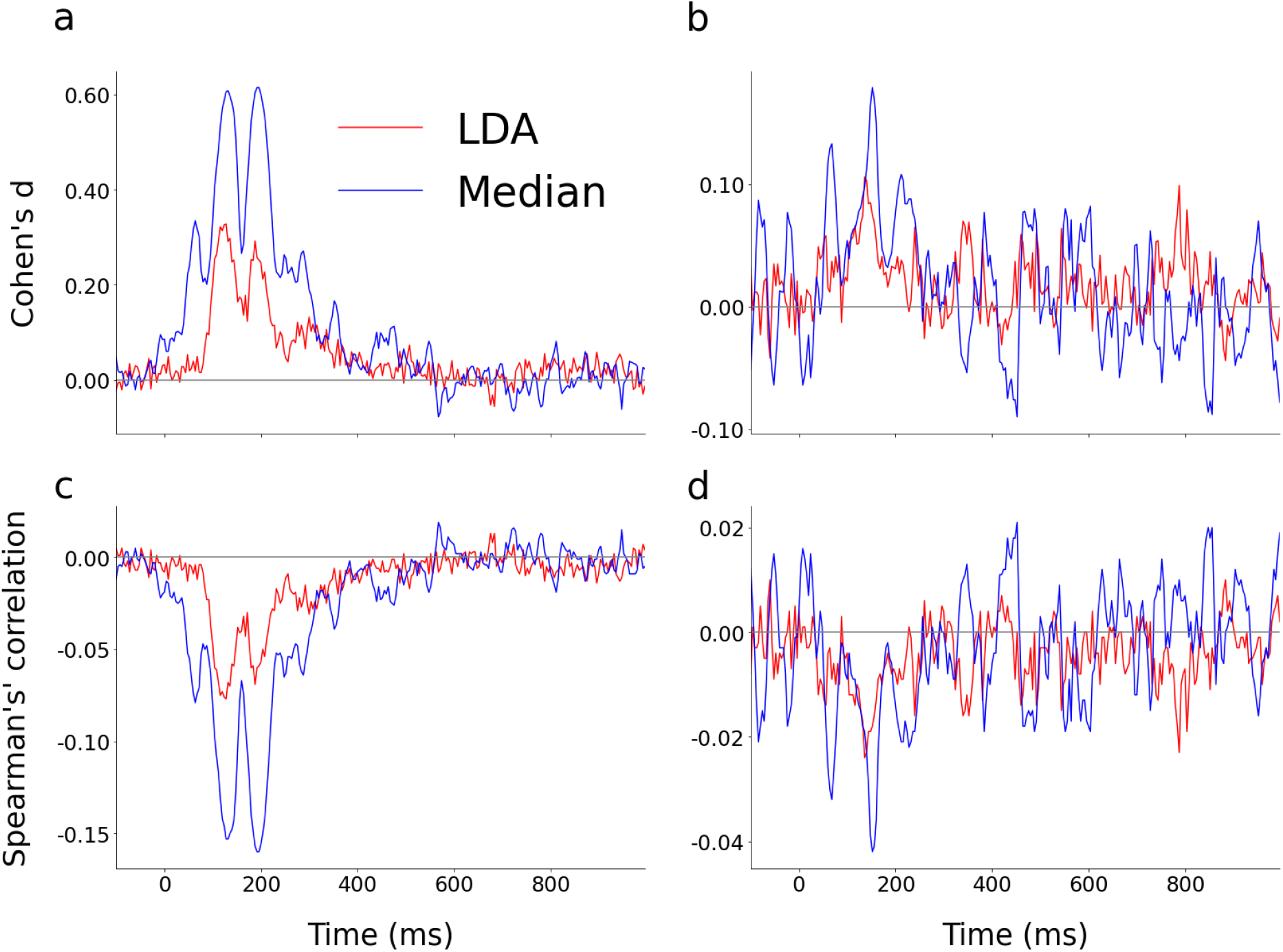
Correspondence between LDA and median accuracy scores, and the first quartile of the principal component distribution. A) Difference between the LDA and median accuracy scores for concepts within the first quartile of the first principal component vs. the rest. B) Same as in panel A, but for the second principal component. D) Spearman correlation between the dissimilarity vector (computed from the first principal component) and the LDA and median scores. D) Same as in panel C, but for the second principal component.

As a complementary analysis, we separated concepts based on the image statistics descriptors following a clustering procedure (see Methods for further information), obtaining four clusters which are represented in **Figure 6A**. We then obtained the effect size of the accuracy scores (LDA, median) between pairs of concepts that belong to the same cluster vs. those that belong to different clusters. This analysis informs how well the concepts can be categorized solely based on the image descriptors. Again, we observe higher effect sizes for the local model (median) for all clusters (**Figure 6B-6E**), implying more vulnerability against regularities in the image statistics. The Spearman correlation coefficients (computed between the classification accuracies and dissimilarity vectors indicating if the concepts belong to the same cluster or not) shown in **Figures 6F** to **6I** present results consistent with those shown in the other panels.

**Figure 6.**
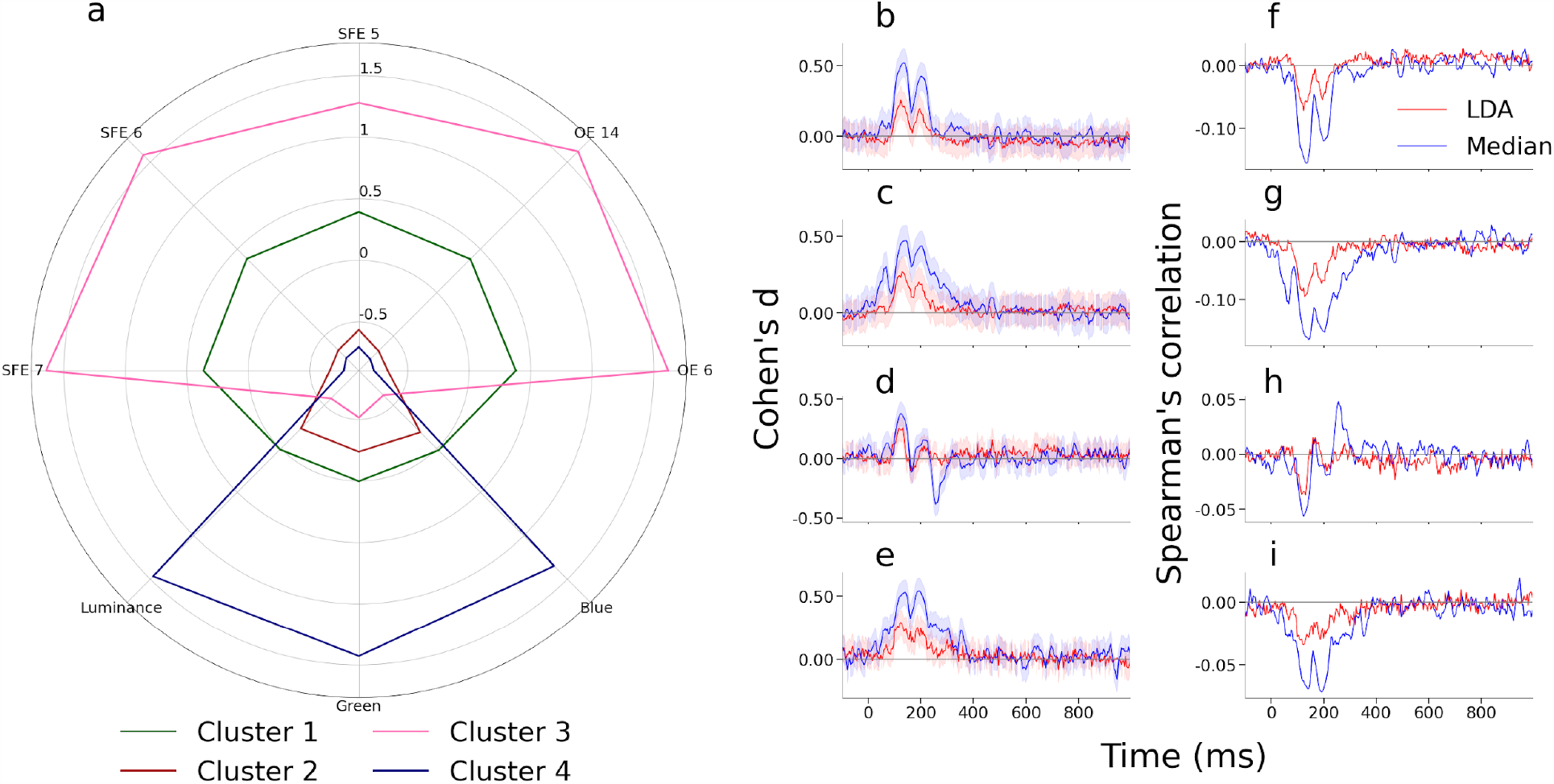
Clustering of concepts based on image statistics descriptors. A) Radar plot representation of the 4 clusters in terms of the original 8 image descriptors. B-E) Difference of the LDA/median accuracy scores between pairs of concepts within the same cluster vs. pairs that belong to different clusters. F-I) Spearman correlation coefficient between LDA/median accuracy scores and the dissimilarity vectors (i.e. whether concepts belong to the same cluster or not).

As a final step in our analysis, we investigated the influence of the image statistics in the categorization of concepts depending on their semantics. We first selected the categories of concepts with more than 1% (>18) of the concepts. From these, we selected a subset given by “human”, “body”, “animal” and “hardware”. These categories were chosen since they contain a large number of concepts and are easily interpretable. We selected all concept pairs within the same category and within different categories, and computed the effect size of the classification scores (LDA and median), with results shown in **Figures 7A** – **7D**. This shows that some concepts in specific categories are easier to discriminate from others, e.g. “animals”, given by the larger classification scores with concepts from other categories. Overall, significant differences were observed for all categories, suggesting they represent a meaningful grouping of the concepts. **Figures 7E** – **7H** give the same information but obtained using Spearman’s correlation coefficient computed between the accuracy scores and the dissimilarity vector, indicating whether the concepts belong to the same or to different categories.

**Figure 7.**
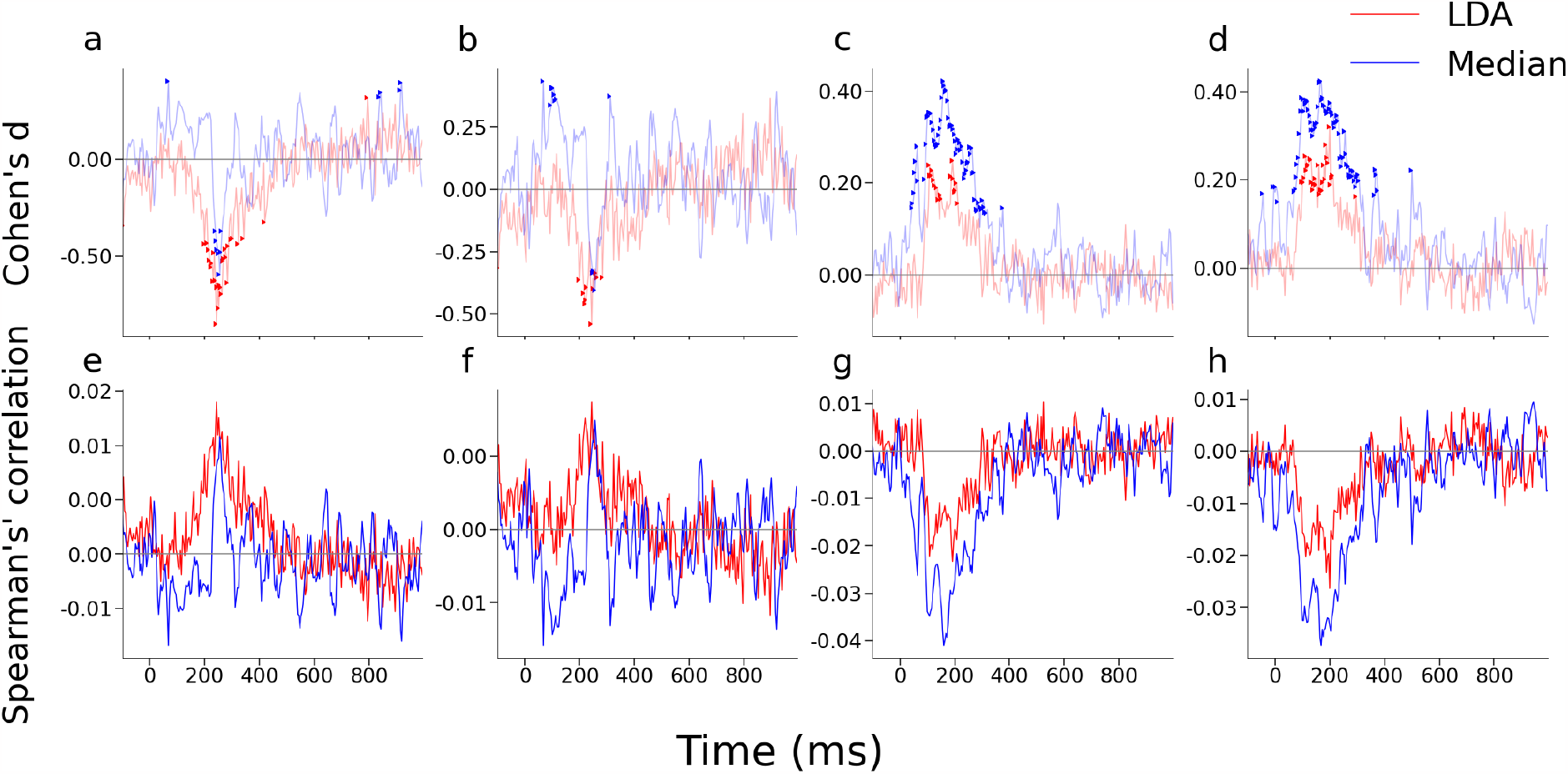
Correspondence between LDA/median classification scores and categories (“human”, “body”, “animal”, and “hardware”). A-D) Effect size of the LDA/median values between concepts within the same category vs. belonging to different categories (from left to right, categories “human”, “body”, “animal”, and “hardware”, respectively. E-H) Spearman correlation coefficient for the LDA/median accuracy scores and the dissimilarity vectors. Points indicate significant results.

After establishing that “human”, “body”, “animal”, and “hardware” are meaningful concept groupings, we implemented an iterative procedure to estimate to which extent the classification by these categories is affected by the natural statistics of the images. The rationale behind the procedure is as follows. For a given category, we first measure the difference between the classification model accuracy (both LDA and median) for all pairs of concepts belonging to this category vs. pairs belonging to different categories (i.e. the selected category and others). Next, we select a concept from the selected category and replace it by another from a different category, but with the most similar image statistic descriptors. If the classification is indeed driven by semantic information, we would expect a decrease in the difference of the accuracies as computed above; however, if the classification is mainly driven by the image statistics, and we replace the concepts with others from different categories but with similar image descriptors, we would not expect a decrease in the difference. We repeated this process iteratively, showing the results in **Figure 8** for categories “human”, “body”, “animal”, and “hardware” (left to right) and LDA/median models (up and bottom panels). We can observe an approximately constant slope for some categories (“animal”, “hardware”) and a clearly decreasing slope for others (“human”, “body”). Interestingly, the categories showing most robustness against degradation of the concepts are those which were the most easily separable (**Figure 7**), suggesting again that a major contributor to the categorization of the images can be found in their low-level statistics. Also consistent with the previous results, the slopes for the median model were smaller than those observed for the LDA model, even though this difference was relatively small.

**Figure 8.**
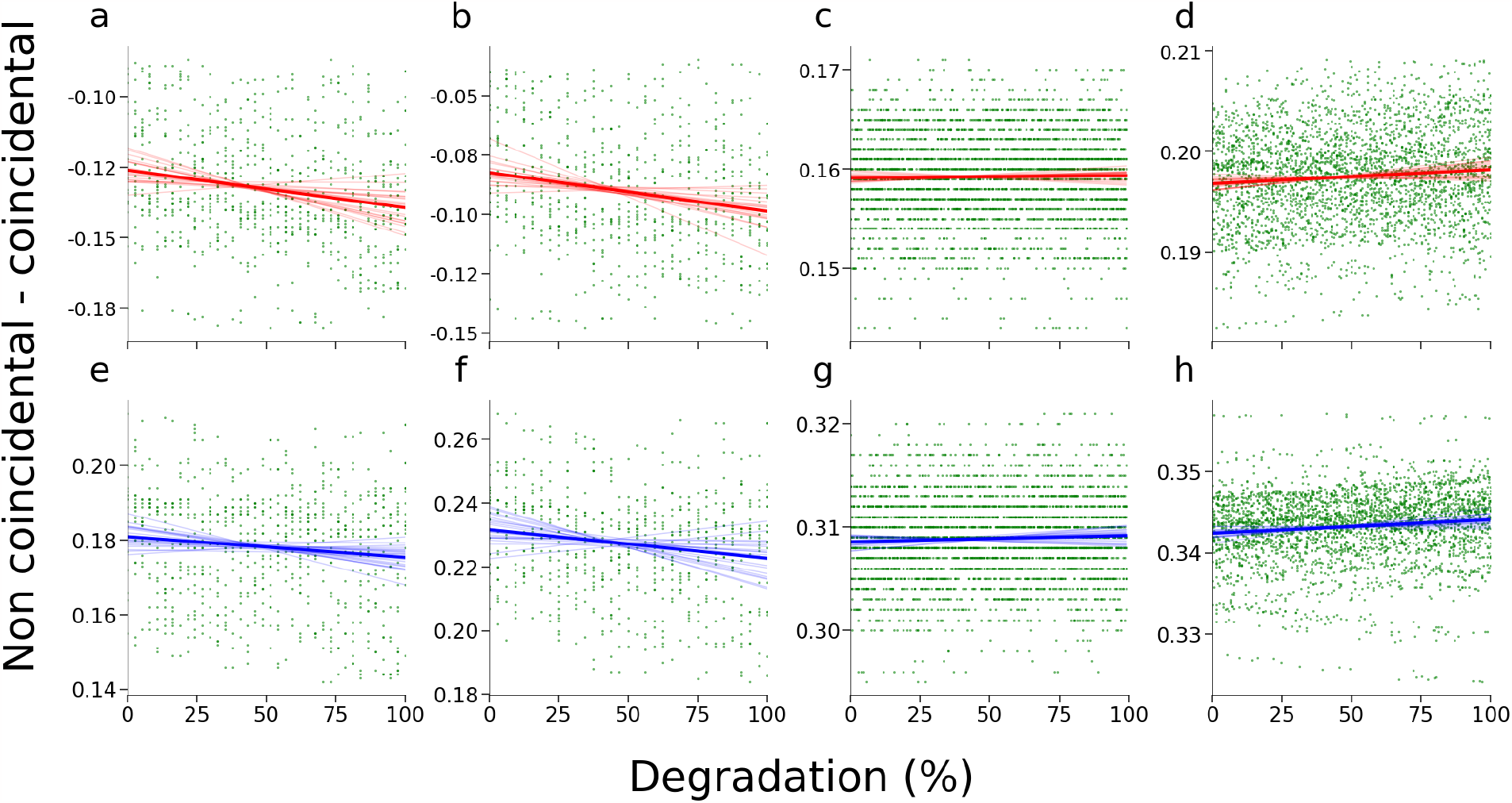
Difference in the LDA / median scores for categories “human”, “body”, “animal”, and “hardware” as a function of the % degradation of the concepts within the category. A) LDA and category “human” (slope = -1e-04; p = 1e-05). B) LDA and category “human” (slope = -1e-04; p = 1e-05). C) LDA and category “animal” (slope = 2e-06; p = 0.24). D) LDA and category “hardware” (slope = 1e-05; p = 7e-05). E) Median and category “human” (slope = -6e-05; p = 5e-04). F) Median and category “body” (slope = -9e-05; p = 5e-04). G) Median and category “animal” (slope = 6e-06; p = 0.03). H) Median and category “hardware” (slope = 2e-05; p = 1e-06).

## Discussion

We investigated the relationship between model complexity and the confounding effect of low-level image descriptors in two different classification tasks based on EEG recordings, both based on images presented in a RSVP paradigm (Grootswagers, Robinson et al. 2019). Our main conclusion is that simple univariate models are capable of outperforming distributed multivariate models based on global information across a wide range of frequencies, but this might occur at the expense of robustness against features that are not directly implicated with semantics. These observations are relevant to the discussion of several fundamental and practical issues in EEG decoding, which are explored below.

In recent years, considerable advances were published in the decoding of semantic information from neuroimaging data across techniques, spanning multiple spatial and temporal resolutions (Robinson, Quek et al. 2023). The use of EEG is especially attractive to resolve the dynamics of object recognition and identification, given its comparatively high temporal resolution (Carlson, Tovar et al. 2013, Cichy, Pantazis et al. 2014, Kaneshiro, Perreau Guimaraes et al. 2015, Cichy, Khosla et al. 2016, Cichy and Teng 2017, Contini, Wardle et al. 2017, Cichy, Kriegeskorte et al. 2019, Kong, Kaneshiro et al. 2020). However, the classification of images based on neuroimaging recordings also presents serious interpretational challenges. Such is the case of the low-level information that confounds image categorization, which can be encoded in features such as luminance, contrast, and many other descriptors computed from the frequency representation of the image. Recently, Harrison demonstrated that image concepts in the THINGS database are separable only based on their luminance and contrast, and that these two features suffice to classify images into concepts with above chance accuracy (Harrison 2022). Besides preparing experimental stimuli which are identical in terms of their low-level information (which may present problems of its own) (Bracci, Ritchie et al. 2017), other solutions include disrupting the high-level category information and assessing the performance of classifiers to distinguish data, thus showing that at least some low-level information tends to contribute to the overall classification accuracy (Long, Stormer et al. 2017). The IMAGE database (Hebart, Contier et al. 2023), as well as other similar efforts (Mehrer, Spoerer et al. 2021), attempt to solve the issue by increasing the variance of the low-level image descriptors by accumulating a much larger and heterogeneous set of images (Grootswagers & Robinson 2021).

The application of increasingly sophisticated machine learning algorithms further complicates these issues, as more complex models are capable of picking small but systematic differences that might not directly inform the meaning of the objects present in the image (Grootswagers & Robinson 2021). In particular, limits to model interpretability hinder the task of detecting the confounding effect of low-level image statistics (Linardatos, Papastefanopoulos et al. 2020). This problem is compounded with limitations intrinsic to the EEG technique, where electrode signals capture a summation of large numbers of neuroelectric sources, and source localization can be challenging and ambiguous (Michel and Murray 2012). In comparison to fMRI, where a large body of knowledge pinpoints the cortical location of low, middle and high-level visual information, EEG lacks this capacity, only providing an estimate of low vs. high-level information in the timing of the components that comprise the measured response (Contini, Wardle et al. 2017, Kong, Kaneshiro et al. 2020, Robinson, Quek et al. 2023). Thus, low-level information (e.g. stimulus position, spatial frequency, orientation, color) appears represented in early signal components (around 50 ms) (Carlson, Hogendoorn et al. 2011, Ramkumar, Jas et al. 2013, Cichy, Ramirez et al. 2015, Robinson, Venkatesh et al. 2017, Blom, Feuerriegel et al. 2020, Teichmann, Quek et al. 2020, Rosenthal, Singh et al. 2022) while semantic information is generally encoded in a later (>100 ms) phase of the response (Carlson, Tovar et al. 2013, Cichy, Pantazis et al. 2014, Grootswagers, Robinson et al. 2019). This raises the possibility of an interaction between model complexity and neurobiological informativeness, where simple models perform comparatively well, but at the expense of representing low-level image information instead of semantic content. This scenario is consistent with an initial phase where low-level information is processed locally at specialized visual cortical regions, giving rise to a later and distributed response which contains information relevant for stimulus categorization (Robinson, Quek et al. 2023). Of note, in this scenario the model performance might not be a robust indicator of sensitivity to semantic content, especially in cases of low signal-to-noise ratio.

Our work builds upon previous efforts to establish the weight of low-level image descriptors in the classification accuracy in object recognition tasks. For instance, Harrison recently showed that high classification accuracy can be obtained using a model trained using low-level image features only (Harrison 2022). However, this does not guarantee that EEG-based decoding operates based on the same information, and thus that EEG-based decoding performance can be explained from these results. In contrast, our analyses provide more compelling evidence supporting this possibility. As shown in **Figures 4** to **6**, not only the image descriptors are correlated with EEG responses, but EEG decoding performance also presents considerable differences between image pairs with similar low-level features vs. pairs that are more dissimilar in this regard. Moreover, as shown in **Figure 7**, it is possible to iteratively replace a member of a category by another with different meaning but similar image statistics while preserving the within group classifier accuracy. This directly implies that, at least in some cases, EEG-based classification accuracy is not driven by semantic similarity but by image similarity instead. Importantly, this occurs for the most populated categories, suggesting that increasing the number of samples (and thus the variance of the corresponding image descriptors) might not suffice to mitigate this confound (Grootswagers & Robinson 2021, Hebart, Contier et al. 2023).

To adequately assess whether EEG signals can encode semantic information about images, several modifications could be introduced to the RSVP experimental paradigm adopted by Grootswagers and colleagues, as well as to the methodology used to train and evaluate the classifiers (Grootswagers, Robinson et al. 2019). An straightforward improvement would be the use of images with homogeneous low-level descriptors, which could be facilitated by the use of simpler backgrounds, e.g. with single colors or regular patterns, such as checkerboards (Bracci, Ritchie et al. 2017). However, to the extent that semantics cannot be fully dissociated from at least part of the low-level information present in the image, this approach risks hindering the identification of the objects by the participants. Another possibility is the exhaustive exploration of multiple (potentially nested) categories, including supersets such as “animals”, “tools” or “body parts”, but also others with more abstract interpretations, such as “animated” or “humanness”, among other possibilities (Bracci, Ritchie et al. 2019, Ritchie, Zeman et al. 2021). At the methodological level, our work suggests that multivariate models based on distributed information across the scalp may be more robust against the confounds of low-level statistics, even if their overall classification performance is lower in comparison to simpler univariate models. A reasonable guiding principle would be the prioritization of models with clearer neurobiological interpretations, whose features can be mapped to the known spatio-temporal dynamics of object recognition in the visual system; in the case of EEG, this could correspond to the timing of the peak decoding performance (Hebart & Baker et al. 2018, Carlson TA 2020).

The interpretation of EEG decoding models could also be benefited by their application to other sensory domains, as well as to the implementation of transfer learning between modalities (Hebart, Contier et al. 2023, Watanabe, Miyoshi et al. 2023). In particular, the latter possibility is interesting since the low-level regularities present in images do not represent a confound for the categorization of sounds, and vice-versa; however, this task is likely more difficult than the classification of unimodal percepts, since it may require the involvement of higher order trans-modal cortical regions, potentially implicating the consolidation of conscious perception, and thus rendering the RSVP paradigm suboptimal for this purpose (Del Cul, Baillet et al. 2007). Another interesting possibility is the study of sensory modalities where the informative vs. non-informative parts of the stimuli can be precisely dissected beforehand. A prime example is the study of olfaction, which is frequently relegated in humans, but has recently been tackled using scalp EEG recordings (Kato, Okumura et al. 2022). The recognition of odorant molecules is based on their chemical composition, mainly the presence of certain functional groups and their location within the molecule and relative to other similar groups (Keller, Gerkin et al. 2017). By exploring the response to molecules with different degrees of structural similarity, it could be possible to dissociate the contribution of the odorant functional groups from other factors. In contrast, this dissociation is more complicated in vision since a certain concept can be presented in multiple ways to a viewer depending on factors such as lightning, position in the field of view, occlusions by other objects, etc. (Robinson, Quek et al. 2023).

In conclusion, our work used multiple approaches to validate previous reports suggesting a confounding effect of low-level image description in the THINGS database. Since there are sufficient reasons to suspect that similar confounds are manifest in other databases of EEG responses to visual stimuli, we investigated the robustness against these factors in terms of model complexity, finding a dissociation between the former and model performance. Future studies could attempt to mitigate these issues by crafting novel sets of stimuli, but also by focusing on the nature of the trained models and their neurobiological interpretability, as well as by investigating sensory modalities where the informativeness of different stimuli features can be more precisely determined.

## References

Blom, T., D. Feuerriegel, P. Johnson, S. Bode and H. Hogendoorn (2020). “Predictions drive neural representations of visual events ahead of incoming sensory information.” Proc Natl Acad Sci U S A 117(13): 7510–7515.

Bonner, M. F. and R. A. Epstein (2021). “Object representations in the human brain reflect the co-occurrence statistics of vision and language.” Nat Commun 12(1): 4081.

Bracci, S., J. B. Ritchie and H. O. de Beeck (2017). “On the partnership between neural representations of object categories and visual features in the ventral visual pathway.” Neuropsychologia 105: 153–164.

Bracci, S., J. B. Ritchie, I. Kalfas and H. P. Op de Beeck (2019). “The Ventral Visual Pathway Represents Animal Appearance over Animacy, Unlike Human Behavior and Deep Neural Networks.” J Neurosci 39(33): 6513–6525. Campbell,

F. W. and J. G. Robson (1968). “Application of Fourier analysis to the visibility of gratings.” J Physiol 197(3): 551–566.

Carlson, T., D. A. Tovar, A. Alink and N. Kriegeskorte (2013). “Representational dynamics of object vision: the first 1000 ms.” J Vis 13(10).

Carlson TA G. T., Robinson AK (2020). An introduction to time-resolved decoding analysis for M/EEG. The Cognitive Neurosciences. G. M. D Poeppel, MS Gazzaniga. Cambridge, MIT Press.

Carlson, T. A., H. Hogendoorn, R. Kanai, J. Mesik and J. Turret (2011). “High temporal resolution decoding of object position and category.” J Vis 11(10).

Cichy, R. M., A. Khosla, D. Pantazis, A. Torralba and A. Oliva (2016). “Comparison of deep neural networks to spatio-temporal cortical dynamics of human visual object recognition reveals hierarchical correspondence.” Sci Rep 6: 27755.

Cichy, R. M., N. Kriegeskorte, K. M. Jozwik, J. J. F. van den Bosch and I. Charest (2019). “The spatiotemporal neural dynamics underlying perceived similarity for real-world objects.” Neuroimage 194: 12–24.

Cichy, R. M., D. Pantazis and A. Oliva (2014). “Resolving human object recognition in space and time.” Nat Neurosci 17(3): 455–462.

Cichy, R. M., F. M. Ramirez and D. Pantazis (2015). “Can visual information encoded in cortical columns be decoded from magnetoencephalography data in humans?” Neuroimage 121: 193–204.

Cichy, R. M. and S. Teng (2017). “Resolving the neural dynamics of visual and auditory scene processing in the human brain: a methodological approach.” Philos Trans R Soc Lond B Biol Sci 372(1714).

Contini, E. W., S. G. Wardle and T. A. Carlson (2017). “Decoding the time-course of object recognition in the human brain: From visual features to categorical decisions.” Neuropsychologia 105: 165–176.

de Bruine, G., A. Vredeveldt and P. J. van Koppen (2018). “Cross-cultural differences in object recognition: Comparing asylum seekers from Sub-Saharan Africa and a matched Western European control group.” Appl Cogn Psychol 32(4): 463–473.

Del Cul, A., S. Baillet and S. Dehaene (2007). “Brain dynamics underlying the nonlinear threshold for access to consciousness.” PLoS Biol 5(10): e260.

Geisler, W. S. (2008). “Visual perception and the statistical properties of natural scenes.” Annu Rev Psychol 59: 167–192.

Gifford, A. T., K. Dwivedi, G. Roig and R. M. Cichy (2022). “A large and rich EEG dataset for modeling human visual object recognition.” Neuroimage 264: 119754.

Grootswagers, T. and A. K. Robinson (2021). “Overfitting the Literature to One Set of Stimuli and Data.” Front Hum Neurosci 15: 682661.

Grootswagers, T., A. K. Robinson and T. A. Carlson (2019). “The representational dynamics of visual objects in rapid serial visual processing streams.” Neuroimage 188: 668–679.

Grootswagers, T., I. Zhou, A. K. Robinson, M. N. Hebart and T. A. Carlson (2022). “Human EEG recordings for 1,854 concepts presented in rapid serial visual presentation streams.” Sci Data 9(1): 3.

Harrison, W. J. (2022). “Luminance and Contrast of Images in the THINGS Database.” Perception 51(4): 244–262.

Hebart, M. N. and C. I. Baker (2018). “Deconstructing multivariate decoding for the study of brain function.” Neuroimage 180(Pt A): 4–18.

Hebart, M. N., O. Contier, L. Teichmann, A. H. Rockter, C. Y. Zheng, A. Kidder, A. Corriveau, M. Vaziri-Pashkam and C. I. Baker (2023). “THINGS-data, a multimodal collection of large-scale datasets for investigating object representations in human brain and behavior.” Elife 12.

Huth, A. G., S. Nishimoto, A. T. Vu and J. L. Gallant (2012). “A continuous semantic space describes the representation of thousands of object and action categories across the human brain.” Neuron 76(6): 1210–1224.

Kaneshiro, B., M. Perreau Guimaraes, H. S. Kim, A. M. Norcia and P. Suppes (2015). “A Representational Similarity Analysis of the Dynamics of Object Processing Using Single-Trial EEG Classification.” PLoS One 10(8): e0135697.

Kato, M., T. Okumura, Y. Tsubo, J. Honda, M. Sugiyama, K. Touhara and M. Okamoto (2022). “Spatiotemporal dynamics of odor representations in the human brain revealed by EEG decoding.” Proc Natl Acad Sci U S A 119(21): e2114966119.

Keller, A., R. C. Gerkin, Y. Guan, A. Dhurandhar, G. Turu, B. Szalai, J. D. Mainland, Y. Ihara, C. W. Yu, R. Wolfinger, C. Vens, L. Schietgat, K. De Grave, R. Norel, D. O. P. Consortium, G. Stolovitzky, G. A. Cecchi, L. B. Vosshall and P. Meyer (2017). “Predicting human olfactory perception from chemical features of odor molecules.” Science 355(6327): 820–826.

Kong, N. C. L., B. Kaneshiro, D. L. K. Yamins and A. M. Norcia (2020). “Time-resolved correspondences between deep neural network layers and EEG measurements in object processing.” Vision Res 172: 27–45.

Kriegeskorte, N., M. Mur and P. Bandettini (2008). “Representational similarity analysis - connecting the branches of systems neuroscience.” Front Syst Neurosci 2: 4.

Kuwabara, M. and L. B. Smith (2016). “Cultural differences in visual object recognition in 3-year-old children.” J Exp Child Psychol 147: 22–38.

Linardatos, P., V. Papastefanopoulos and S. Kotsiantis (2020). “Explainable AI: A Review of Machine Learning Interpretability Methods.” Entropy (Basel) 23(1).

Logothetis, N. K. and D. L. Sheinberg (1996). “Visual object recognition.” Annu Rev Neurosci 19: 577–621.

Long, B., V. S. Stormer and G. A. Alvarez (2017). “Mid-level perceptual features contain early cues to animacy.” J Vis 17(6): 20.

Masarwa, S., O. Kreichman and S. Gilaie-Dotan (2022). “Larger images are better remembered during naturalistic encoding.” Proc Natl Acad Sci U S A 119(4).

Mehrer, J., C. J. Spoerer, E. C. Jones, N. Kriegeskorte and T. C. Kietzmann (2021). “An ecologically motivated image dataset for deep learning yields better models of human vision.” Proc Natl Acad Sci U S A 118(8).

Michel, C. M. and M. M. Murray (2012). “Towards the utilization of EEG as a brain imaging tool.” Neuroimage 61(2): 371–385.

O’Toole, A. J., F. Jiang, H. Abdi and J. V. Haxby (2005). “Partially distributed representations of objects and faces in ventral temporal cortex.” J Cogn Neurosci 17(4): 580–590.

Ramkumar, P., M. Jas, S. Pannasch, R. Hari and L. Parkkonen (2013). “Feature-specific information processing precedes concerted activation in human visual cortex.” J Neurosci 33(18): 7691–7699.

Ritchie, J. B., A. A. Zeman, J. Bosmans, S. Sun, K. Verhaegen and H. P. Op de Beeck (2021). “Untangling the Animacy Organization of Occipitotemporal Cortex.” J Neurosci 41(33): 7103–7119.

Robinson, A. K., G. L. Quek and T. A. Carlson (2023). “Visual Representations: Insights from Neural Decoding.” Annu Rev Vis Sci 9: 313–335.

Robinson, A. K., P. Venkatesh, M. J. Boring, M. J. Tarr, P. Grover and M. Behrmann (2017). “Very high density EEG elucidates spatiotemporal aspects of early visual processing.” Sci Rep 7(1): 16248.

Rosenthal, I. A., S. R. Singh, K. L. Hermann, D. Pantazis and B. R. Conway (2022). “Color Space Geometry Uncovered with Magnetoencephalography.” Curr Biol 32(7): 1670–1674.

Teichmann, L., G. L. Quek, A. K. Robinson, T. Grootswagers, T. A. Carlson and A. N. Rich (2020). “The Influence of Object-Color Knowledge on Emerging Object Representations in the Brain.” J Neurosci 40(35): 6779–6789.

Watanabe, N., K. Miyoshi, K. Jimura, D. Shimane, R. Keerativittayayut, K. Nakahara and M. Takeda (2023). “Multimodal deep neural decoding reveals highly resolved spatiotemporal profile of visual object representation in humans.” Neuroimage 275: 120164.

